# Nod-like receptors are critical for gut-brain axis signaling

**DOI:** 10.1101/647032

**Authors:** Matteo M. Pusceddu, Mariana Barboza, Melinda Schneider, Patricia Stokes, Jessica A. Sladek, Cristina Torres-Fuentes, Lily R. Goldfild, Shane E. Gillis, Ingrid Brust-Mascher, Gonzalo Rabasa, Kyle A. Wong, Carlito Lebrilla, Mariana X. Byndloss, Charles Maisonneuve, Andreas J. Bäumler, Dana J. Philpott, Richard Ferrero, Kim E. Barrett, Colin Reardon, Mélanie G. Gareau

## Abstract

Gut-brain axis signaling is critical for maintaining health and homeostasis. Stressful life events can impact gut-brain signaling, leading to altered mood, cognition and intestinal dysfunction. Here we identify nucleotide binding oligomerization domain (Nod)-like receptors (NLR), Nod1 and Nod2, as novel regulators for gut-brain signaling. NLR are innate immune pattern recognition receptors expressed in the gut and brain, important in the regulation of gastrointestinal (GI) physiology. We found that mice deficient in both Nod1 and Nod2 (NodDKO) demonstrate signs of stress-induced anxiety, cognitive impairment and depression in the context of a hyperactive hypothalamic-pituitary-adrenal axis. These deficits were coupled with impairments in the serotonergic pathway in the brain, decreased hippocampal neurogenesis, and reduced neural activation. In addition, NodDKO mice had increased GI permeability and altered serotonin signaling in the gut following exposure to acute stress. Administration of the selective serotonin reuptake inhibitor, fluoxetine, abrogated behavioral impairments and restored serotonin signaling. We also identified that intestinal epithelial cell-specific deletion of Nod1 (VilCre^+^Nod1^f/f^), but not Nod2, increased susceptibility to stress-induced anxiety-like behavior and cognitive impairment following exposure to stress. Together these data suggest that intestinal epithelial NLR are novel modulators of gut-brain communication and may serve as potential novel therapeutic targets for the treatment of gut-brain disorders.

## INTRODUCTION

The hypothalamic-pituitary-adrenal (HPA) axis is a critical regulator of the stress response. It is one of the main pathways through which the gut-brain axis signals, with stress negatively impacting intestinal function, including permeability, motility (1) and visceral sensitivity (2). Stress can also induce relapse in patients with gastrointestinal (GI) disease, such as inflammatory bowel disease (IBD) (3) and irritable bowel syndrome (IBS) (4). Stressful life events also represent risks factors for the development of psychiatric disorders, including anxiety and major depressive disorder (5). Additionally, there is significant genetic pleiotropy for psychiatric and immune system disorders, suggesting an underlying common pathway of regulation (6). While the precise mechanisms for this susceptibility to stress remain to be fully understood, it is thought to involve a complex interplay between environmental, biological, and genetic risk factors.

The immune system plays a pivotal role in brain function and stress responses (7, 8). Mice deficient in T- and B-cells display anxiety-like behavior and cognitive deficits compared to wild type (WT) controls (9). More recently, mice deficient for the innate immune pattern recognition receptor (PRR), peptidoglycan (PGN) recognition protein (PGLYRP) 2 gene, showed alterations in social behavior in a sex dependent manner (10). Together, these findings suggest a role for adaptive and innate immune pathways in regulating gut-brain communication. Nucleotide binding oligomerization domain (Nod)-like receptors (NLR), Nod1 and Nod2, are intracellular PRRs that recognize specific moieties of PGN and are important in maintaining gut homeostasis by eliciting protective immune responses to bacteria (11, 12). In addition to their well-established expression in the periphery, including in intestinal epithelial cells (IEC) and mucosal immune cells (13, 14), Nod1 and Nod2 are expressed in several brain areas, including the hippocampus and diverse cell types, such as pyramidal neurons, astrocytes, and microglia (10). Despite this evidence of central nervous system (CNS) expression, the precise role of Nod1 and Nod2 in the brain remains to be fully elucidated.

Given their physiological distribution, we hypothesized that NLR may represent a novel potential therapeutic target for the treatment of stress-related disorders and a novel modulator of gut-brain communication. Using mice deficient in both Nod1 and Nod2 (NodDKO), we assessed behavior and brain function as well as intestinal physiology in the context of acute psychological stress.

## METHODS

### Mice

Adult (6-8 week) male and female mice (Jackson Labs; bred in-house) were used in the study. Nod1/Nod2 double knockout (NodDKO; C57BL/6 background) mice and WT (C57BL/6) controls were maintained at UC Davis and in-bred for at least 11 generations (15). Nod1 floxed mice were created using a targeting construct for *Nod1* generated using C57BL/6-derived bacterial artificial chromosome clones. The construct was designed with *loxP* sites flanking exons 2 and 3 of *Nod1* and an EGFP kanamycin/neomycin cassette was introduced for selection in *Escherichia coli* and embryonic stem (ES) cells, respectively. The targeting construct was verified by sequencing and electroporated into C57BL/6 ES cells. G418-resistant ES cell clones were picked and expanded. DNA was extracted and screened for targeting by PCR. ES cell clones that amplified a product of the expected size were further characterized by Southern blot analysis to confirm targeting at both the 5’ and 3’ ends of the targeting construct. Correctly targeted ES cell clones were karyotyped and two independently targeted clones with a normal chromosome count were prepared for microinjection into blastocysts. Chimeric mice were prepared by microinjecting gene targeted ES cells into recipient blastocysts (Balb/c) and transferred to pseudo-pregnant recipients. The targeted ES cells were derived from C57BL/6 mice. Male chimeric mice were mated with WT C57BL/6 females to obtain germline transmission of the targeted allele. Animals that were heterozygous for the targeted allele (as determined by coat color) were bred to generate *Nod1*^f/f^ animals which were bred and maintained under specific-pathogen-free (SPF) conditions in the Monash Medical Centre Animal Facilities. Generation of floxed mice was performed according to the guidelines of Monash University’s Institutional Biosafety and Animal Ethics Committees (Suppl Fig 6). Nod2^f/f^ mice were created as described elsewhere (16). Once generated, both Nod1^f/f^ and Nod2^f/f^ mice were bred at UC Davis where all experiments were performed.

Nod1^f/f^ and Nod2^f/f^ mice were crossed with IEC-specific Cre-expressing (VilCre) mice (Jackson Labs; bred in-house). Cages consisted of either VilCre^-^Nod1^f/f^ and VilCre^+^Nod1^f/f^ or VilCre^-^ Nod2^f/f^ and VilCre^+^Nod2^f/f^ genotypes.

Mice were housed in cages lined with chip bedding and had free access to food and water throughout the study. The vivarium lighting schedule allowed 12 hours of light and 12 hours of darkness each day with temperature maintained at 21 ± 1 °C. All behavioral tests were performed between 9am and 6pm and all procedures and protocols were reviewed and approved by the Institutional Animal Care and Use Committee at the University of California, Davis (IACUC protocol #20072).

### Co-housing

Co-housing was performed in a subset of WT and NodDKO mice starting at post-natal day 21. Animals, paired by age and sex, were housed in the same cage for 3-4 weeks.

### Fluoxetine treatment

Fluoxetine hydrochloride (18 mg/Kg per day; Sigma-Aldrich) was administered *ad libitum* in the drinking water in opaque bottles to protect it from light. The drinking water containing fluoxetine was changed every 3 days to prevent any possible degradation. Control animals received plain drinking water as vehicle. Animals were treated immediately starting after weaning for a period of 28 days. On day 28, animals underwent the one-day behavioral battery testing (Fig 4A).

### WAS

Mice were placed on small platforms (inverted 50ml beakers) in a clean, standard housing cage filled with approximately 2 cm of room temperature water for 1 hour. Water avoidance stress (WAS) is a well-established model for inducing stress in rodents (17). After this exposure to stress, mice were returned to their home cage and allowed to rest for 5-10 minutes prior to experimentation or euthanasia (Fig 1A).

**Figure 1.**
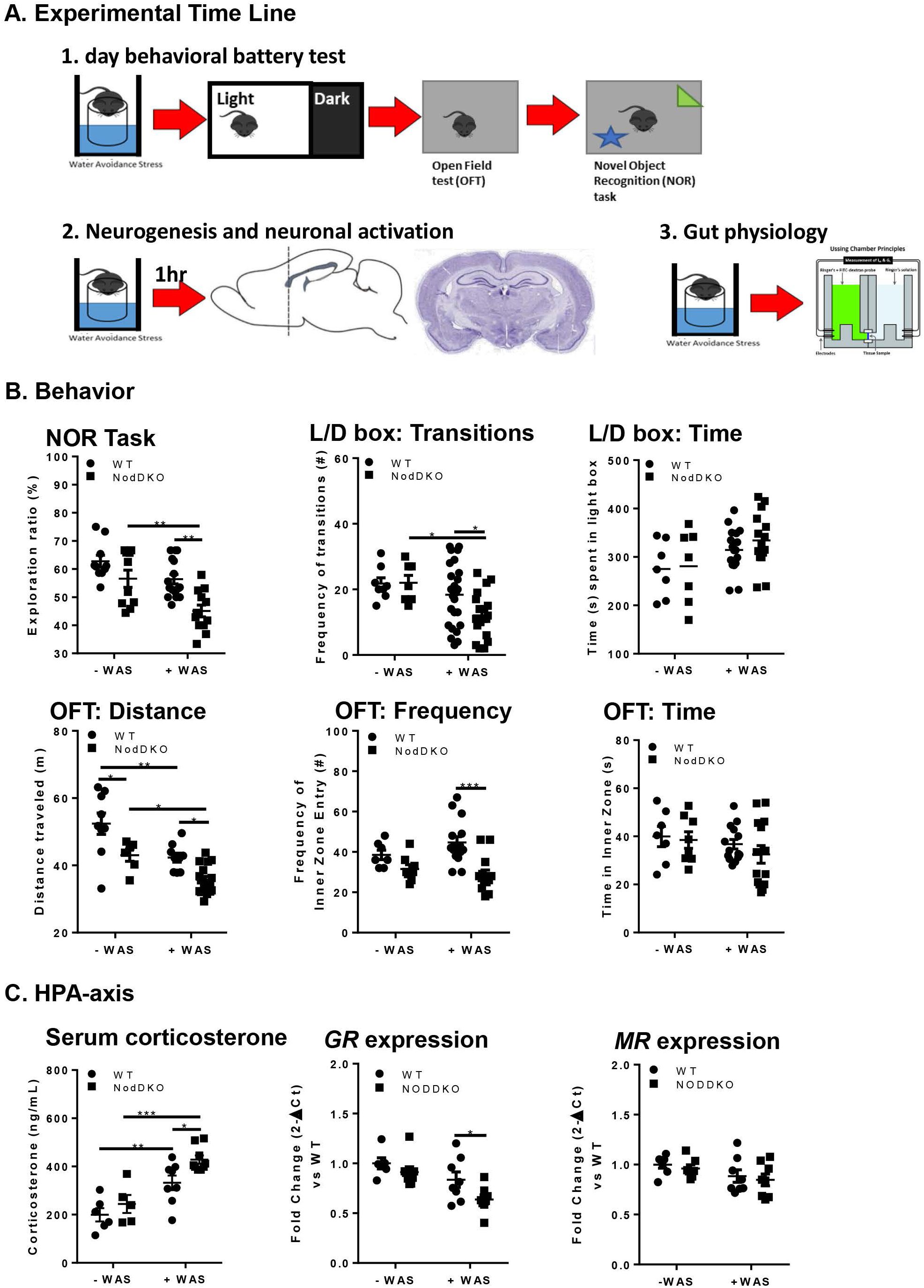
NodDKO(+WAS) mice display behavioral deficits and HPA-axis hyperactivation. (**A**) Experimental time line: adult C57BL/6 male and female WT and NodDKO mice underwent: 1) 1 day behavioral battery testing; or 2) perfusion for hippocampal neurogenesis and neural activation analysis; or 3) gut physiology assessed by Ussing chambers. (**B**) Behavior: novel object recognition (NOR) task, light/dark (L/D) box and open field test (OFT) (N = 10-16); (**C**) HPA-axis: serum corticosterone levels (N = 6-9), hippocampal glucocorticoid (*GR*) and mineralcorticoid (*MR*) receptor mRNA expression levels (N = 6-8) following exposure to 1h of water avoidance stress (WAS). Data are presented as mean ± SEM. (*P < 0.05; **P < 0.01; ***P < 0.001, 2-way ANOVA).

### L/D Box Test

To measure anxiety-like behavior, the light/dark (L/D) box test was used as described previously (17). Briefly, mice were placed in a box with a light (2/3) and a dark (1/3) compartment for 10 minutes, during which behavior of the mouse was video-recorded. These videos were later scored by an observer blinded to treatment paradigms for the proportion of time spent by the mouse in the lit portion of the box, with lower values ascribed to anxiety-like behavior (18). In addition, the number of times the mouse transitioned from the dark compartment to the light compartment of the box were quantified to indicate the mouse’s activity and exploratory level.

### OFT

Animal behavior for the open field test (OFT) was recorded for 10 minutes in the novel arena during acclimation for the novel object recognition (NOR) task. Both locomotor activity and anxiety-like behavior were determined as quantification of total distance moved, time spent in the inner zone, and frequency of inner zone entries as analyzed by a digital tracking system (Ethovision).

### NOR Task

The NOR task was performed as previously described (9, 17). Briefly, mice were subjected to 10 min of acclimation to the novel arena (30 × 30 cm white plexiglass box) followed by 5 min of habituation to two identical objects (training phase). After a rest and recovery period (45 minutes) one of the two original objects was replaced with a novel object and interactions were monitored for an additional 5 min (testing phase). Direct contacts with the objects, including any contact with nose or paw, or an approach within 2 cm, were recorded and scored using Ethovision (XT 8.5, Noldus, Leesburg, VA) and verified manually. Any close contact in which the animal was not directly interacting with the object but rather exploring the chamber were not interpreted as direct contact with the object and omitted. Results are expressed as a ratio quantifying preference for exploration of a novel object rather than a familiar object during the testing phase. An exploration ratio of greater than 50% indicates that the mouse investigated the novel object more than the familiar object, indicating memory recall for the latter (17). Mice that did not pass the training phase (less than 40% or greater than 60% exploration ratio [suggesting bias towards one object]) were omitted.

### FST

The forced swim test (FST) was used to assess depressive-like behavior. Briefly, mice were placed individually in Pyrex cylinders (40 cm tall × 20 cm in diameter) filled with water (24.5 °C) to a 30 cm depth for 6 minutes. The last 4 minutes of the test were analyzed using a tracking system (Ethovision). During the behavioral analysis, mobility time, intended as any movements other than those necessary to balance the body and keep the head above the water, was measured.

### Immunofluorescence

Brains (WT[+WAS] and NodDKO[+WAS]) were collected following anesthesia (5% isoflurane) and transcardial perfusion with 4% PFA. Brains were post-fixed overnight at 4°C and placed in 30% (w/v) sucrose for 4 days before embedding in optimal cutting temperature (OCT) medium in ice-cold isopentane followed by storage at −80°C. Samples were cut using a cryostat (Leica Microsystem, Germany) into 20 µm-thick coronal sections. Sections were serially collected and stored at −20 °C until use. Immunofluorescence was performed as previously described (19). Briefly, sections underwent an antigen retrieval step using citrate buffer (10 mM, pH 6.0, 1h, 95°C). After blocking in 5% BSA normal goat serum (1h, room temperature), samples were incubated with primary antibody overnight (16h, 4°C). The primary antibodies used were: rabbit Anti-c-Fos (2250S, Cell Signaling, Danvers, MA), guinea pig anti-DCX [(AB2253, Millipore, Burlington, MA)], and rabbit anti-Ki67 (LS-C141898, Lifespan Biosciences, Seattle, WA). Slides were washed (3 × 5mins) and incubated with appropriately labeled secondary antibodies (1 h at room temperature), then washed and mounted in Prolong diamond (Invitrogen, Carlsbad, CA). The secondary antibodies used were Alexa 647 goat anti-rabbit (ab150155, Abcam, Cambridge, UK) for both c-Fos and Ki67, and Alexa 555 goat anti-guinea pig for DCX (A21435, Invitrogen, Carlsbad, CA). DAPI was used as a nuclear stain.

### Image analysis and cell quantification

Confocal imaging was performed on a Leica SP8 STED 3X microscope. Immunofluorescent Z-stack images with a 1.04 µm step size were collected using a 20x objective. Systematic random sampling was used for the dorsal hippocampus by counting the cells in both hemispheres of each section in 1:6 series (120 µm apart). Every second section, for a total of three sections, was used either for doublecortin (DCX)/ Ki67 co-staining [dentate gyrus (DG) area] or for c-Fos staining [cornus ammonium (CA)3 and DG areas]. Cell quantification was performed using the image processing software package Imaris x64_8.2.1. All cell numbers are expressed as an average of 3 sections per animal. The dorsal hippocampus was defined as AP −0.94 to −2.30 according to the Paxinos and Franklin atlas of the mouse brain (20).

### Serum corticosterone

Blood samples were collected via cardiac puncture following CO_2_ exposure. Blood was centrifuged and serum aspirated and stored at −80°C. Corticosterone levels were assayed using a commercially-available enzyme immune assay (EIA) kit (Corticosterone EIA Kit, ADI-900-097, Enzo Life Sciences) according to the manufacturer’s instructions. Absorbance was read with a multi-mode plate reader (Synergy H1, BioTek Instruments, Inc.) at 405 nm.

### Quantitative PCR

Tissues [hippocampus, prefrontal cortex (PFC), colon and ileum] were collected and frozen at −80°C before homogenization in Trizol (Invitrogen). RNA was isolated according to the manufacturer’s protocol (Invitrogen), treated with DNase 1 (Invitrogen) and transcribed into cDNA (BioRad, iSCRIPT cDNA Synthesis kit; Proflex PCR System, Applied Biosystem). qPCR was performed using SYBR green and β-actin used as a housekeeping gene on a QuantStudio 6 Flex Real time PCR machine (Applied Biosystem). Data were presented as ΔΔCT. All primer sequences used in this study are shown in Table 1.

### Brain serotonin (5-HT) quantification

The hippocampus and brain stem were freshly dissected and stored at −80°C until analysis. Tissues were homogenized in 20 mM HEPES buffer (pH 7.5) containing 0.25M sucrose and a cocktail of protease inhibitors (Calbiochem, Burlington, MA) followed by ultracentrifugation at 100,000g for 45 min. Membrane-free tissue supernatants (100 µl) were transferred to 96-well plates for analysis. Liquid chromatography-mass spectrometry (LC/MS) analysis was performed on an Agilent Infinity 1290 ultra-high-performance LC coupled to a triple quadropole MS. Chromatographic separation was carried out on an Agilent Pursuit 3 Pentafluorophenyl (PFP) stationary phase (2 × 150 mm, 3 mm) column. The mobile phases were 100% methanol (B) or water with 0.25% formic acid (A). The analytical gradient was as follows: 2%-60% B in 2 min, 100% B from 2-4 min, 2% B for 4 min. Flow rate was 300 µl/min. Samples were held at 4°C in the autosampler, and the column was operated at 40°C. The MS was operated in positive and selected reaction monitoring (SRM) mode. A 5-HT standard was used for optimization and to generate the calibration curve for quantification. Data acquisition and processing was done using Mass Hunter Qualitative and Quantitative Analysis (Version B.06.00) and Quantitative Analysis (Version B.08.00).

### Serum tryptophan (Trp) quantification

Serum proteins were removed by ultrafiltration using Amicon Ultra Centrifugal Filters (molecular weight cutoff 3KDa). Briefly, the upper chambers of the centrifugal devices were rinsed with water (400 µL) twice and centrifuged at 10,000 g for 5 min before individual mouse serum samples (50 µl) were loaded. Water (150 µl) was added to the samples and they were centrifuged at 10,000 g for 5 min. Protein-free serum flow-throughs were collected in 1.5 mL tubes and aliquots (50 µl) were transferred into vials for MS-analysis as described above.

### Ussing chambers

Segments of GI tract (distal ileum and proximal colon) were excised and cut along the mesenteric border and mounted in Ussing chambers (Physiologic Instruments, San Diego, CA), exposing 0.1 cm^2^ of tissue area to 4 ml of circulating oxygenated Ringer’s buffer maintained at 37°C. The buffer consisted of (in mM) of: 115 NaCl, 1.25 CaCl_2_, 1.2 MgCl_2_, 2.0 KH_2_PO_4_ and 25 NaHCO_3_ at pH 7.35±0.02. Additionally, glucose (10 mM) was added to the serosal buffer as a source of energy, which was balanced osmotically by mannitol (10 mM) in the mucosal buffer. Agar–salt bridges were used to monitor potential differences across the tissue and to inject the required short-circuit current (Isc) to maintain the potential difference at zero as registered by an automated voltage clamp. A computer connected to the voltage clamp system recorded Isc and voltage continuously and analyzed using acquisition software (Acquire and Analyze; Physiologic Instruments). Baseline Isc values were obtained at equilibrium, approximately 15 min after the tissues were mounted. Isc, an indicator of active ion transport, was expressed in μA/cm^2^. Conductance (G) was used to assess tight junction permeability and mucosal to serosal flux of 4KDa FITC-labeled dextran (Sigma) over time (sampled every 30 minutes for 2 hours) was used to assess macromolecular permeability. After completion of the FITC flux measurements, tissues were treated with forskolin (FSK, 20 *μ*M) to assess viability.

### Study Design

Three sets of animals were used for three parallel experiments.

#### 1) Behavioral testing

Mice (±WAS) were exposed to the L/D box test, the OFT, and NOR task (Fig 1A) followed by a 2 hours rest and finally the FST (in a subset of mice; Fig 4A). Animals were habituated to the testing room by allowing them to acclimate in their home-cages in the testing room for at least 30 min prior to testing. Once the behavioral tests were completed, mice were euthanized by CO_2_ inhalation followed by cervical dislocation, and tissues were collected for further analysis.

#### 2) Brain imaging

Mice exposed to 1 hour of WAS were left undisturbed in their home-cages for 1 hour followed by measurement of c-FOS expression levels in the hippocampus (21). Animals were then perfused as described above.

#### 3) Ussing chambers

Mice (±WAS) were euthanized by CO_2_ inhalation followed by cervical dislocation and both ileum and colon were collected to measure ion transport and permeability in Ussing chambers.

### Statistical analysis

Results are expressed as means ± standard error (SEM). Unpaired Student’s t-test or two-way ANOVA followed by Tukey’s post hoc test were performed as appropriate using Prism 7 GraphPad (San Diego, CA). A *P*-value of less than 0.05 was selected as the threshold of statistical significance.

## RESULTS

### NodDKO mice display stress-induced behavioral deficits

To study the role of NLR in regulating stress responses, behavior was assessed in NodDKO mice. NOR task was used to assess recognition memory. While baseline cognitive functions in NodDKO mice were intact, as determined by the ratio of interactions between a known and a novel object, exposure to a single session of acute WAS for 1h led to a cognitive deficit compared to (WT[+WAS]) mice and NodDKO(-WAS) control mice (Fig 1B). In the L/D box test, used to assess anxiety-like behavior, NodDKO(-WAS) mice did not demonstrate evidence of baseline anxiety-like behavior, as indicated by total number of transitions between the L/D boxes and, time spent in the light box compared to WT(-WAS) controls (Fig 1B). In contrast, NodDKO(+WAS) mice displayed a significant decrease in the number of transitions between the L/D compartments compared to WT(+WAS) mice without an impact on time spent in the light box, indicative of anxiety-like behavior. The OFT was used to measure exploratory behavior and general well-being along with serving as a secondary assessment of anxiety-like behavior. NodDKO(+WAS) mice displayed significantly reduced frequency of entries into the inner zone in the OFT compared to WT mice, without a difference in the total time spent in the inner zone (Fig 1B). Exploratory behavior, as determined by total distance travelled in the OFT, was significantly reduced in NodDKO(-WAS) mice compared to WT, which was further decreased in NodDKO(+WAS). Stress also caused decreased exploratory behavior in WT(+WAS) mice. These findings suggest that deficiency in NLR increases susceptibility to acute stress, profoundly impacting behavior.

Cognitive function and anxiety-like behaviors were similar in male and female mice for both WT(±WAS) and NodDKO(±WAS) groups, therefore, all subsequent data are derived from combined sexes, which were equally distributed (Suppl Fig 1A).

### NodDKO mice display hyperactivation of the HPA-axis

Dysregulation of the HPA-axis is associated with the development of anxiety-like behavior (22) and cognitive impairments (23). Given that exposure to WAS impaired behavior in NodDKO mice, we measured serum corticosterone levels and expression of glucocorticoid receptors (GR) and mineralocorticoid receptors (MR) in the hippocampus, which serves as the primary site for feedback inhibition of HPA-axis signaling. As expected, WAS increased the concentration of serum corticosterone in WT mice, with this stress response being further increased in NodDKO(+WAS) compared to WT(+WAS) mice (Fig 1C). As basal corticosterone serum concentrations were similar between WT(-WAS) and NodDKO(-WAS) mice, these data suggest NodDKO mice are susceptible to stress-induced hyperactivation of the HPA axis (Fig 1C).

Corticosterone induces physiological and behavioral effects by binding to GR and MR, which are expressed in the hippocampus and serve to limit HPA-axis activation. Decreased GR function and expression are implicated in the stress response and have been associated with depression (24, 25). Hippocampal mRNA expression of *GR* was significantly reduced in NodDKO(+WAS) mice compared to WT(+WAS) mice, whereas no differences in *MR* expression were observed (Fig 1C). Decreases in *GR* and *MR* in NodDKO(+WAS) mice were also seen in the PFC region (Suppl Fig 2B).

### NodDKO mice have impaired stress-induced neural activation

Neural activation is critical for memory formation with acute stress-induced increases in glucocorticoids necessary for hippocampal neuronal survival, memory acquisition, and consolidation (26). To identify the neural circuitry underlying the differential stress susceptibility between NodDKO(+WAS) and WT(+WAS) mice, we measured expression of the immediate early genes (IEGs) *Arc* and c-Fos, indicative of neural activation (27, 28). *Arc* mRNA expression was significantly increased in the hippocampus of WT(+WAS) but not NodDKO(+WAS) mice compared with WT(-WAS) or NodDKO(-WAS) mice respectively (Fig 2A). This was specific for the hippocampus, with no changes in *Arc* seen in the PFC (Suppl Fig 2B). These findings suggest that Nod1/2 are required for stress-induced activation of hippocampal neurons and subsequent memory consolidation. In contrast, brain-derived neurotrophic factor (*BDNF*), which is important for maintaining neuronal survival, was not impacted in either the hippocampus or PFC in NodDKO(+WAS) mice (Suppl Fig 2A-B).

**Figure 2.**
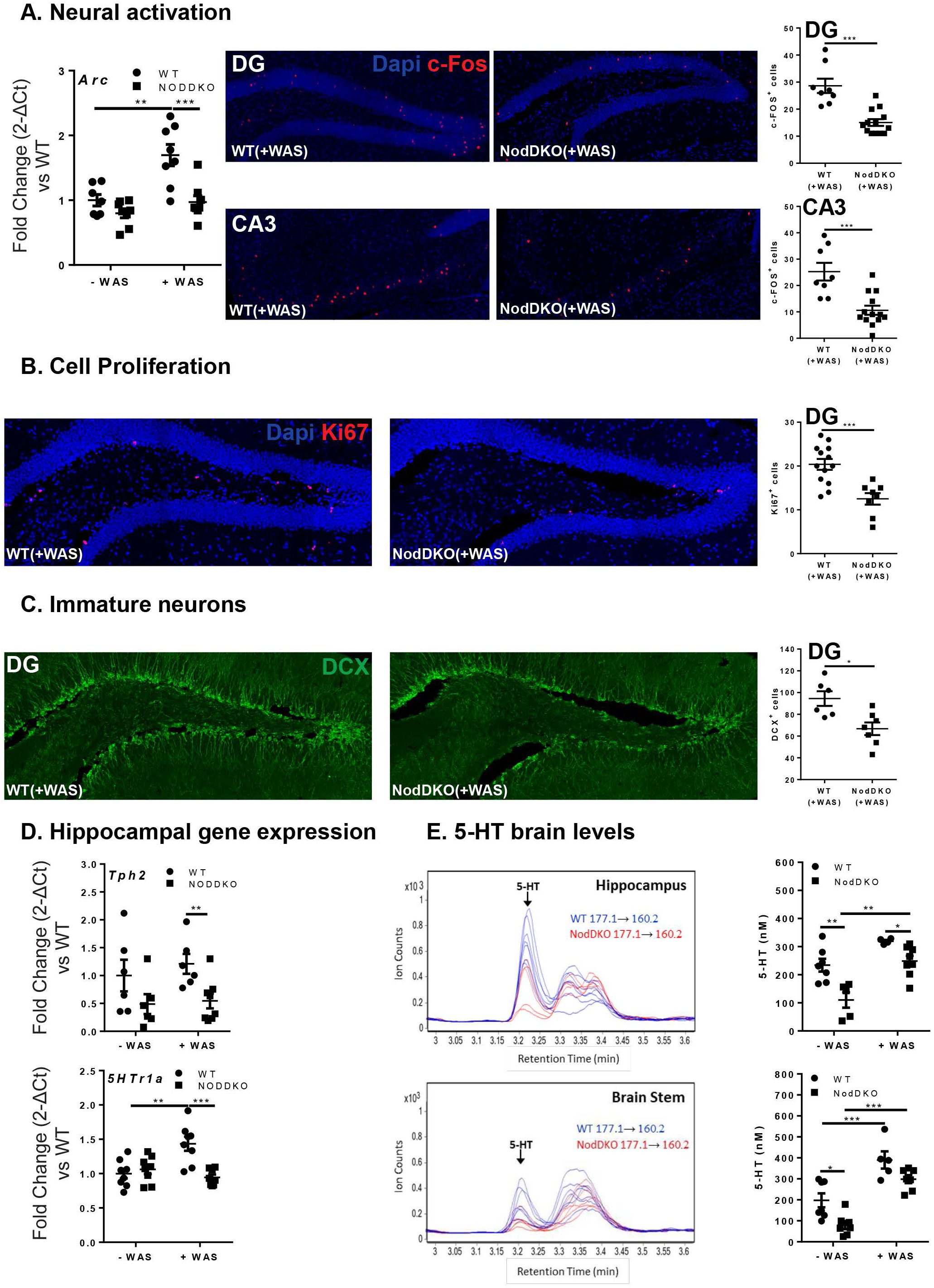
NodDKO(+WAS) mice display decreased neural activation, impaired neurogenesis and down-regulation of the 5-HT system. (**A**) *Arc* mRNA expression levels and c-Fos^+^ cells in the dentate gyrus (DG) (N = 8-13) and the cornus ammonis (CA) 3 (N = 8-13). Representative photographs of c-Fos immunofluorescence. (**B**) Ki67^+^ cells (N = 8-13) and representative photographs of Ki67 immunofluorescence in the DG. (**C**) DCX^+^ cells (N = 6-7) and representative photographs of DCX immunofluorescence in the DG. (**D**) *Tph2* and *5HTr1a* hippocampal mRNA expression levels (N = 6-8); (**E**) serotonin (5-HT) protein levels in the hippocampus (N = 5-8) and brain stem (N = 5-7) detected by LC/MS and multiple reaction monitoring chromatogram of 5-HT in cytoplasmic extracts of both the hippocampus and brain stem: m/z 177.1 correspond to 5-HT precursor ion mass and m/z 160.2 correspond to fragment ion mass. Data are presented as mean ± SEM. (*P < 0.05; **P < 0.01; ***P < 0.001, 2-way ANOVA and unpaired Student’s t-test).

Neuronal activation was further assessed by quantification of c-Fos-positive cells in the DG and CA3 regions of the dorsal hippocampus by confocal microscopy in mice exposed to WAS. Given that the peak of c-Fos expression occurs 1-3 h following exposure an acute stimulus (21), c-Fos positive cells were enumerated in hippocampal sections collected at 1 h post-WAS. The total number of c-Fos-positive cells, as quantified using Imaris, was significantly lower in NodDKO(+WAS) mice compared to WT(+WAS) in both the DG and CA3 region of the dorsal hippocampus (Fig 2A). Taken together, these results suggest that neuronal activation following WAS is impaired in NodDKO(+WAS) mice, leading to lack of consolidation of hippocampal-dependent memory, and subsequently leading to cognitive deficits compared to WT(+WAS) mice.

### NodDKO(+WAS) mice exhibit reduced neural proliferation and decreased neurogenesis in the dorsal hippocampus

Adult hippocampal neurogenesis is thought to play an important role in the stress response (29), consolidation of memory (30), and regulation of mood (31). Given the impaired recognition memory and increased anxiety-like behavior observed in NodDKO(+WAS) mice, we assessed if hippocampal neuronal proliferation and neurogenesis were impacted by Nod1/2 deficiency following WAS. Proliferation of new neurons was measured by staining for the proliferation marker Ki67 and neurogenesis was assessed by quantifying the number of immature neurons stained for DCX in the dorsal hippocampus. Confocal image analysis revealed significantly fewer Ki67-positive cells (Fig 2B) and lower numbers of DCX-positive cells in the DG of NodDKO(+WAS) mice compared to WT(+WAS) controls (Fig 2C). These findings suggest Nod1/2 may regulate stress-induced hippocampal neurogenesis.

### NodDKO mice exhibit down-regulation of the serotonergic system in the brain

Changes in the central 5-HT system have been shown to regulate adult hippocampal neurogenesis in rodents (32, 33). To investigate the potential molecular mechanisms underlying the behavioral deficits observed in NodDKO(+WAS) mice, we first assessed components of the 5-HT signaling pathway by PCR. In the hippocampus, expression of tryptophan hydroxylase (*Tph*)*2*, the rate limiting enzyme for 5-HT synthesis, and the 5-HT receptor (*5HTr*)*1a*, which regulates 5-HT release in different brain areas including the hippocampus (34), were significantly reduced in NodDKO(+WAS) mice when compared to WT(+WAS) controls (Fig 2D). Expression of *5HTr2c* in contrast was not impacted in either brain region, hippocampus and PFC (Suppl Fig 2A-B).

To determine the impact of reduced Tph2 expression on 5-HT in the hippocampus and the brain stem concentrations, LC/MS was used. The brain stem was assessed as it represents the primary site of 5-HT synthesis in the brain (35). Quantification of 5-HT revealed significantly reduced baseline concentrations in the hippocampus and brain stem of NodDKO(-WAS) mice compared to WT(-WAS) controls (Fig 2E). 5-HT was increased by WAS in WT (brain stem) and NodDKO (hippocampus and brain stem) mice, with WT(+WAS) having higher hippocampal 5-HT levels compared to NodDKO(+WAS). These data demonstrate that Nod1/2 participate in an important control mechanism that regulates 5-HT concentration in the brain and impairs stress-induced 5-HT receptor expression.

Other neurotransmitters such as gamma-aminobutyric acid (GABA), are involved in the regulation of stress, anxiety, and cognition (36). Overall, no overt alterations were observed in GABA signaling either in the hippocampus or PFC with only *Gabab1b* found to be increased in the PFC of NodDKO(+WAS) mice compared to WT(+WAS) (Suppl Fig 2A-B). These findings suggest a specific interaction between NLR and serotonergic signaling.

### NodDKO mice exhibit stress-induced impairments in intestinal physiology

Psychological stress and the resulting HPA-axis activation can detrimentally impact intestinal physiology, in part by causing elevated intestinal ion transport and increased intestinal permeability (37). Given previous findings identifying a role for Nod1 and Nod2 in regulating intestinal mucosal barrier function (38), we assessed intestinal physiology in ileal and colonic tissues from WT and NodDKO mice using Ussing chambers. Active ion transport was assessed by measuring Isc in ileal and colonic segments, with NodDKO(-WAS) mice displaying normal baseline values compared to WT(-WAS) mice (Fig 3A). Intestinal permeability was assessed by measuring G for tight junction permeability and flux of FITC-labeled dextran (4KDa) for macromolecular permeability. Similarly to ion transport, G and FITC flux were both normal in NodDKO(-WAS) mice compared to WT(-WAS) mice. In contrast, both ion transport (Isc) and permeability (G and FITC-dextran flux) were increased in colon and ileum from NodDKO(+WAS) mice compared to WT(+WAS) controls (Fig 3A) suggesting that the dysregulated HPA-axis in NodDKO mice detrimentally impacts intestinal physiology.

**Figure 3.**
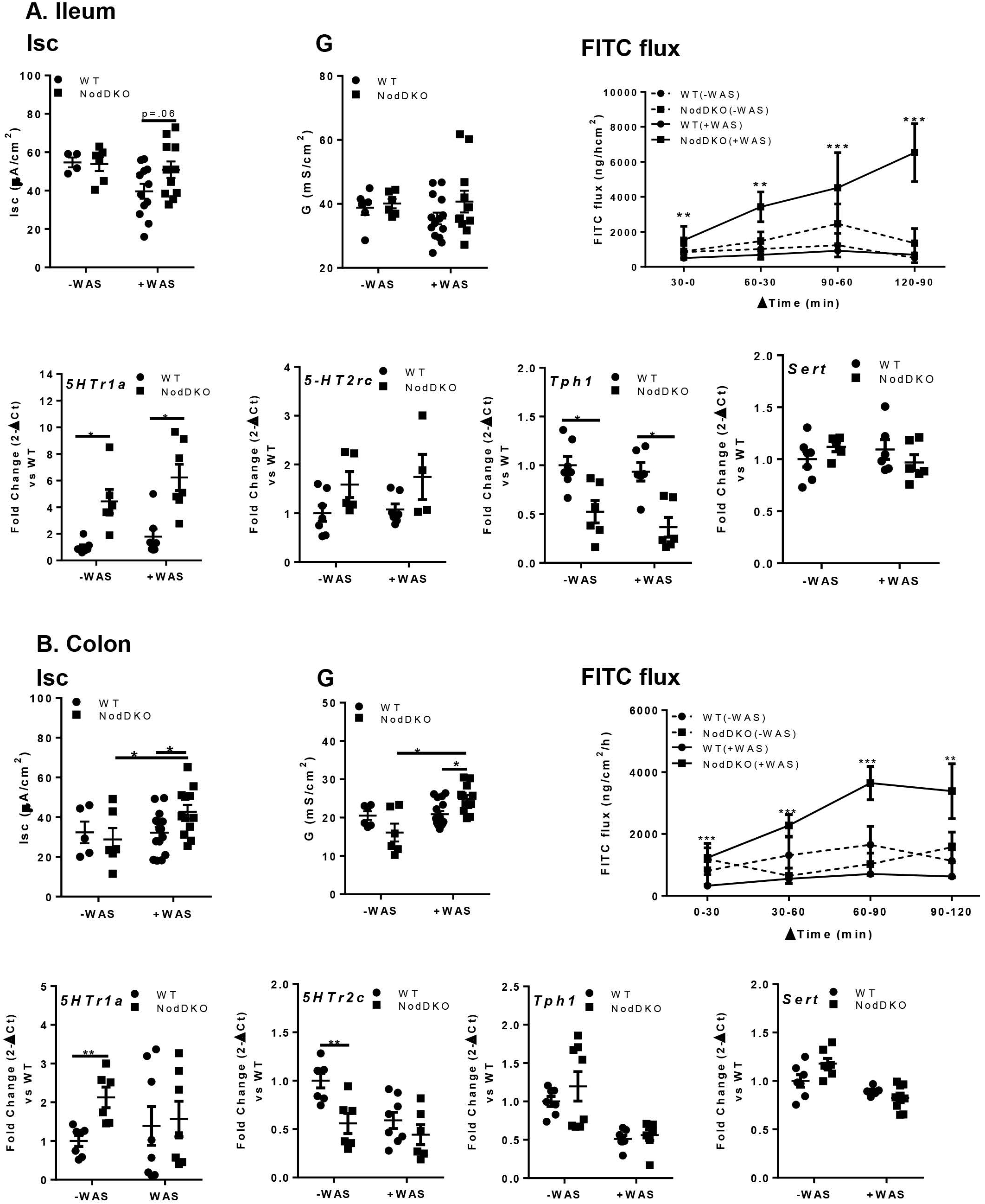
NodDKO(+WAS) mice exhibit increased gastrointestinal permeability and altered 5-HT system. (**A**) Ileum and (**B**) colon basal short circuit current [Isc], basal conductance [G] and FITC dextran flux assessment (N = 11-16) as well as *5HTr1a, 5HTr2c, Tph1* and *SERT* mRNA expression levels (N = 6-8). Data are presented as mean ± SEM. (*P < 0.05; **P < 0.01; ***P < 0.001, unpaired Student’s t-test and 2-way ANOVA).

**Figure 4.**
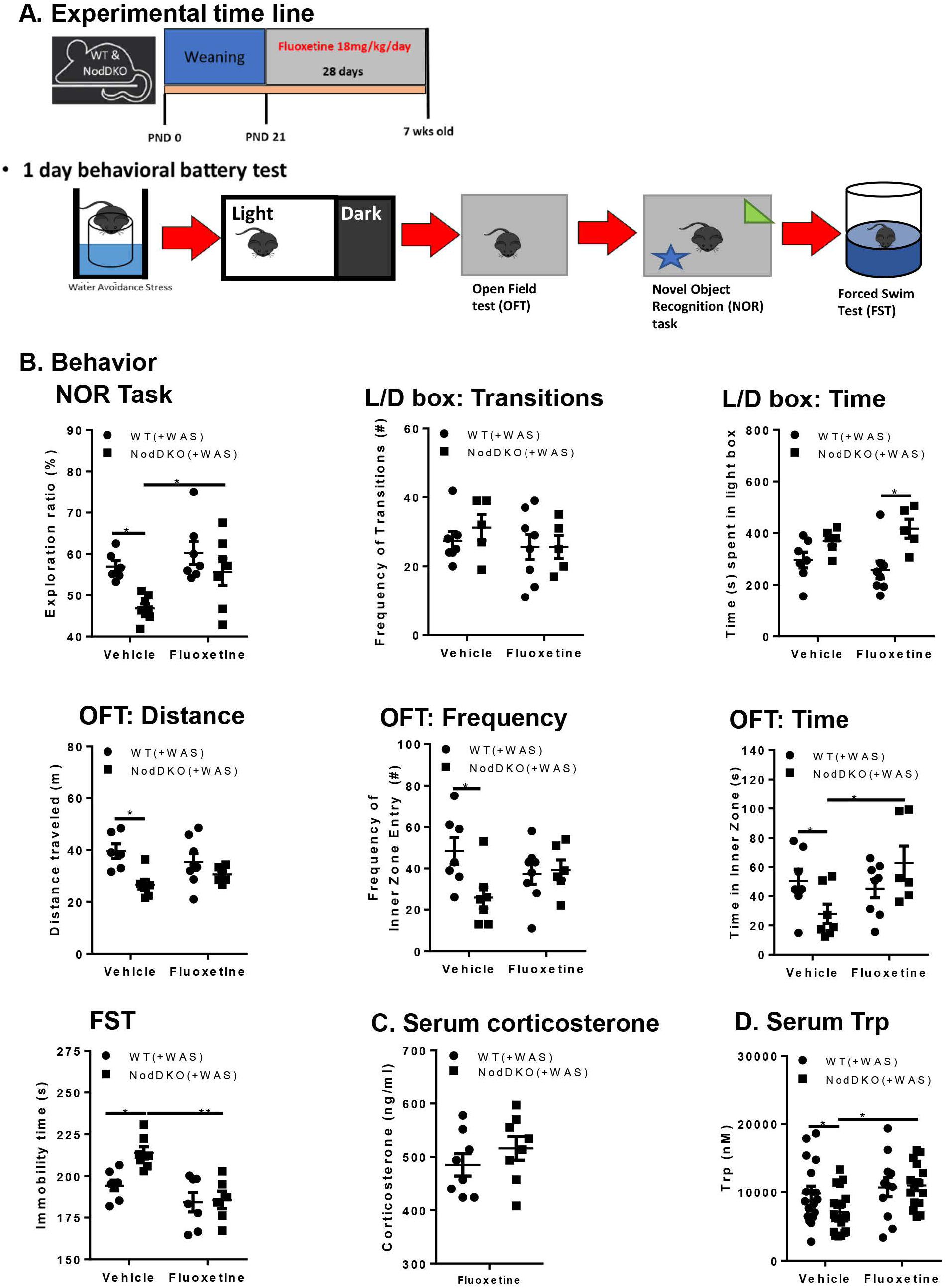
Chronic fluoxetine administration restored behavioral impairments as well as corticosterone and tryptophan serum levels in NodDKO(+WAS) mice. (**A**) Experimental time line showing fluoxetine treatment and the 1 day behavioral battery testing. Fluoxetine effect on (**B**) NOR task, L/D box, OFT and the forced swim test (FST) (N = 6-8); (**C**) Corticosterone (Cort) (N = 8) and (**D**) Tryptophan (Trp) serum levels detected by LC/MS (N = 11-20). Data are presented as mean ± SEM. (*P < 0.05; **P < 0.01; ***P < 0.001, 2-way ANOVA).

Given the important role of 5-HT signaling in regulating gut physiology (39), and our evidence of altered 5-HT signaling in the hippocampus in NodDKO(+WAS) mice, qPCR analysis for 5-HT signaling in ileum and colon were performed. Reduced *Tph1* and increased *5HTr1a* expression were observed in the ileum of NodDKO(+WAS) mice when compared to WT(+WAS) controls (Fig 3A). In the colon, expression of *5HTr1a* and *5HTr2c* were both increased whereas *Tph1* was unaffected (Fig 3B). Expression of the serotonin reuptake transporter (*SERT),* which terminates 5-HT activity, was unaffected in both ileum and colon (Fig 3A, B). Taken together, these findings identify that Nod1 and Nod2 deficiency leads to dysregulated peripheral 5-HT signaling, similar to findings in the hippocampus.

### Stress-induced behavioral impairments in NodDKO mice are restored by chronic fluoxetine administration

The selective serotonin reuptake inhibitors (SSRI), fluoxetine, is one of the most commonly prescribed anti-depressant drugs that ameliorates cognitive impairments and depression both in rodents (40) and humans (41), mainly by acting on the 5-HT system (42, 43). Our observation of reduced 5-HT and 5-HT signaling was down-regulated in NodDKO(+WAS) mice, mice were treated chronically with fluoxetine for 28 days starting at weaning to restore hippocampal 5-HT levels. Cognitive impairments seen in NodDKO(+WAS) mice in the NOR task relative to WT(+WAS) were reversed by fluoxetine administration when compared to vehicle (water)-treated control NodDKO(+WAS) mice (Fig 4B). Fluoxetine administration also improved anxiety-like behavior by increasing the time spent in the L/D box, but not affecting the number of transitions, in NodDKO(+WAS) mice compared to WT(+WAS) fluoxetine-treated mice (Fig 4B). Similarly, deficits in total distance traveled, time spent, and frequency of entries into the inner zone of the OFT in NodDKO(+WAS) mice were reversed by fluoxetine treatment compared to vehicle controls (Fig 4B). Given fluoxetine’s anti-depressant properties, depressive-like behavior was assessed using the FST. The total time spent immobile was increased in NodDKO(+WAS) mice compared to WT(+WAS) mice administered vehicle, suggesting the presence of depressive-like behavior. This was reversed by chronic fluoxetine administration to NodDKO(+WAS) mice (Fig 4B). Chronic fluoxetine administration also abolished the elevation of serum corticosterone levels that had previously been seen (Fig 1C) in NodDKO(+WAS) mice compared to WT(+WAS) mice suggesting the ability of fluoxetine to modulate HPA-axis activation (Fig 4C).

### Nod1/2 deficiency reduces serum tryptophan levels, which can be restored by chronic fluoxetine administration

As the essential amino acid Trp is the sole precursor of peripherally and centrally produced 5-HT (44), and reduced serum Trp levels correlate with alterations in mood and cognition (45, 46), serum Trp concentrations were assessed by LC/MS. Lower serum Trp levels were found in both NodDKO(-WAS) (Suppl Fig 3) and NodDKO(+WAS) than in their control WT(±WAS) counterparts (Fig 4D), suggesting a role for Nod1/2 in regulating Trp transport. Interestingly, chronic fluoxetine treatment restored serum Trp concentration to WT levels in NodDKO(+WAS) mice (Fig 4D).

### Chronic fluoxetine administration did not restore intestinal physiology in NodDKO(+WAS)

Given the impairments in 5-HT signaling found in the ileum and colon of NodDKO(+WAS) mice, we assessed if chronic fluoxetine administration could restore the altered intestinal physiology we observed in these animals. In contrast to amelioration of behavioral deficits, the elevations in secretory state (Isc) and gut permeability (G and FITC flux) seen in NodDKO(+WAS) in ileum (Suppl Fig 4A) and colon (Suppl Fig 4B) were not reversed by chronic fluoxetine administration. These findings suggest that while there is a role for 5-HT signaling in regulating intestinal physiology in NodDKO(+WAS), the mechanism by which fluoxetine normalizes behavior is likely independent of intestinal physiology.

### Stress-induced behavioral deficits are dependent on intestinal epithelial Nod1 receptors

To determine whether intestinal NLR account for the behavioral and biochemical effects observed in NodDKO mice, we utilized a conditional knockout (cKO) strategy. Behavior and intestinal physiology were assessed in IEC cKO mice (VilCre^+^Nod1^f/f^ and VilCre^+^Nod2^f/f^). VilCre^+^Nod1^f/f^(+WAS) mice demonstrated significantly reduced recognition of the novel object in the NOR task compared to control mice (VilCre^-^Nod1^f/f^[+WAS]; Fig 5A). Furthermore,

**Figure 5.**
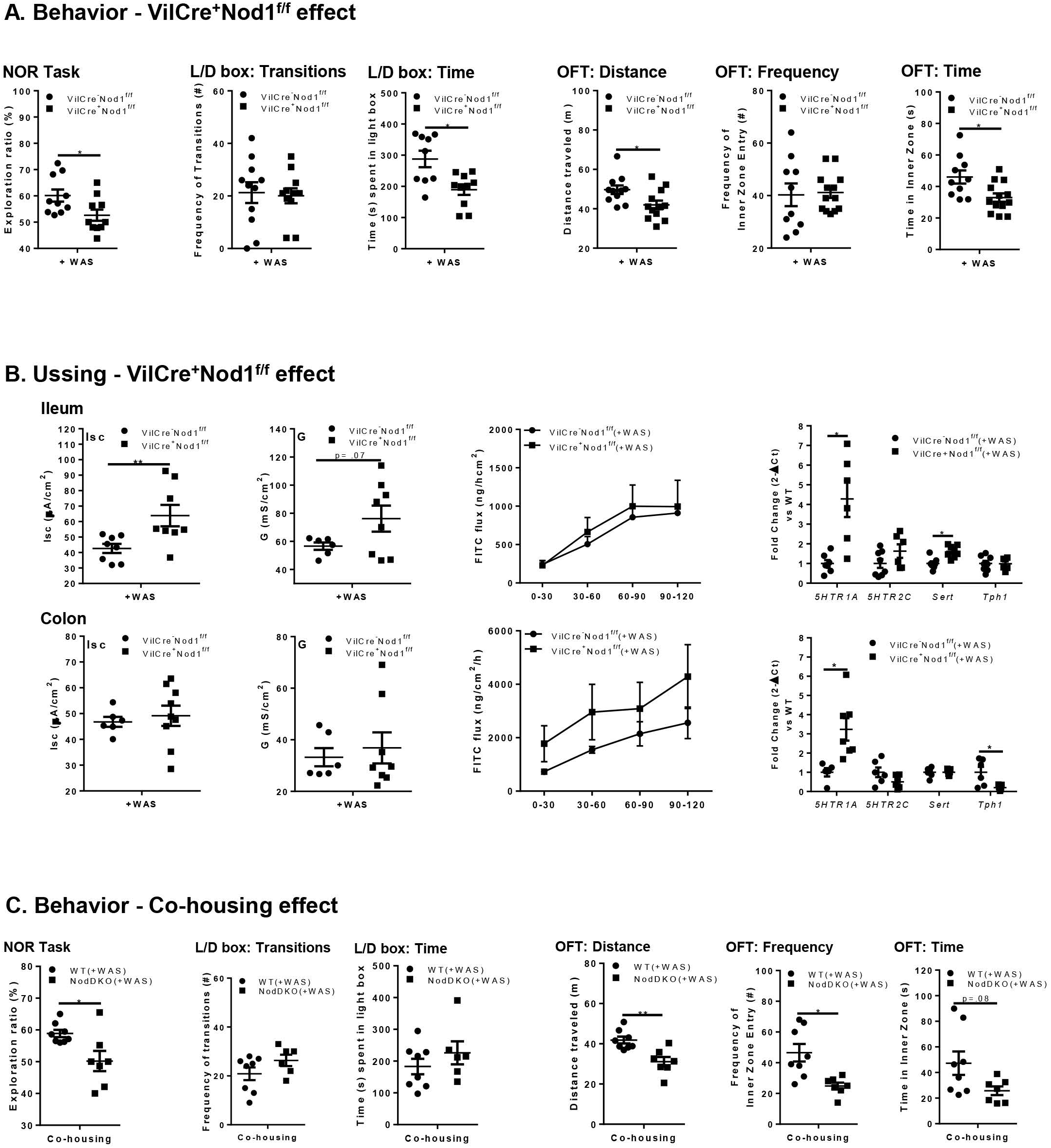
Behavioral deficits in NodDKO(+WAS) mice are dependent on intestinal epithelial Nod1 receptor expression but are not rescued by co-housing. Effect of intestinal Nod1 deletion on (**A**) NOR task, L/D box and OFT (N = 7-12). (**B**) Colon and ileum basal short circuit current [Isc], basal conductance [G] and FITC dextran flux assessment (N = 6-9) as well as *5HTr1a, 5HTr2c, SERT* and *Tph1* mRNA expression levels (N = 6-8). (**C**) Co-housing effect between WT(+WAS) and NodDKO(+WAS) on NOR task, L/D box and OFT (N = 7-12). Data are presented as mean ± SEM. (*P < 0.05; **P < 0.01, unpaired Student’s t-test).

VilCre^+^Nod1^f/f^(+WAS) mice displayed anxiety-like behavior as determined by the L/D box (reduced time spent in the light compartment) and OFT (reduced total distance traveled and time spent into the inner zone) (Fig 5A). In contrast, VilCre^+^Nod2^f/f^(+WAS) mice had normal cognitive function and anxiety-like behavior when compared to littermate controls (Suppl Fig 5). These findings suggest that IEC Nod1 expression can regulate CNS mechanisms and impact behavior.

In the intestinal tract, significantly increased ion transport (Isc) was found in the ileum of VilCre^+^Nod1^f/f^(+WAS) mice compared to their WT control littermates, (Fig 5B). In contrast, no changes in physiology were observed in the colon, with similar ion transport and permeability seen in both WT and VilCre^+^Nod1^f/f^(+WAS) mice. These findings suggest a role for intestinal epithelial Nod1 expression in the regulation of intestinal physiology, while also highlighting an important role for Nod2, given the reduced impacts of the single cKO compared to the NodDKO mice. qPCR analysis for 5-HT signaling in ileum and colon identified increased *5HTr1a* and *SERT* mRNA expression levels in VilCre^+^Nod1^f/f^(+WAS) mice when compared to VilCre^-^ Nod1^f/f^(+WAS) controls (Fig 5B). In the colon, *5HTr1a* was increased whereas *Tph1* mRNA levels were decreased (Fig 5B). Taken together, these findings suggest that IEC Nod1 expression can regulate peripheral 5-HT signaling, but the precise impact depends on the gut segment studied.

### Behavioral deficits in NodDKO(+WAS) mice are not rescued by co-housing

To assess whether behavioral alterations and biochemical changes that occurred in NodDKO(+WAS) mice were a reflection of their genotype or were driven by the microbiota, we co-housed a subset of mice starting at post-natal day 21, pairing age and sex-matched WT and NodDKO mice in the same cage. This co-housing strategy did not prevent development of cognitive deficits and anxiety-like behavior in NodDKO(+WAS) mice compared to cage mate WT(+WAS) control mice (Fig 5C). In the NOR task, the co-housed NodDKO(+WAS) mice showed a lower exploration ratio compared to co-housed WT(+WAS) littermates. With respect to anxiety-like behavior, no differences were observed in the L/D box test, while co-housed NodDKO(+WAS) mice showed evidence of anxiety-like behavior in the OFT as indicated by lower total distance travelled and decreased frequency of inner zone entry (Fig 5C). These findings suggest that the composition of the microbiota does not contribute markedly to the behavioral deficits seen in NodDKO(+WAS) mice, such that abnormal behaviors cannot be transferred to WT(+WAS) mice within the same cage.

## DISCUSSION

Understanding the molecular mechanisms underlying stress susceptibility is key for the identification of novel pharmacological treatments for stress-related gut-brain disorders. Here we demonstrated that mice deficient in Nod1 and Nod2 have altered serotonergic signaling in the gut and brain, which makes them susceptible to hyperactivation of the HPA-axis following exposure to acute psychological stress. This HPA-axis activation leads to GI pathophysiology, cognitive deficits, anxiety-like behavior and depressive-like behavior. Highlighting a previously unappreciated role of Nod1/2 in regulating HPA-axis activation and serotonergic signaling, behavioral deficits (but not abnormalities in GI physiology) in NodDKO(+WAS) mice were normalized by chronic fluoxetine administration. Using a cKO approach, we further identified that mice lacking Nod1 in IEC and exposed to WAS recapitulated the changes in behavior and intestinal physiology seen in NodDKO(+WAS) mice. Deficiency of Nod1 in IEC in these mice resulted in stress-induced impairments in cognitive function and anxiety-like behavior. These effects appear to reflect a specific function of Nod1 in IEC, as there was no impact of loss of IEC Nod2 expression on behavior or GI physiology. Together, these results identify Nod1 as a novel factor that regulates the stress response, 5-HT biosynthesis, and signaling. These results further indicate that Nod1 may contribute to a previously unrecognized signaling pathway in the gut-brain axis.

As NLR are expressed in both the brain and periphery, and function as receptors that detect bacterial PGN, NLR were hypothesized to modulate gut-brain signaling. Highlighting this novel role, mice deficient in Nod1/2 subjected to WAS had pronounced cognitive dysfunction, as well as anxiety-like and depressive-like behaviors, compared to WT(+WAS) controls. These findings complement previous studies by Farzi *et al.*, who showed a role for NLR in regulating sickness behaviors (47). In contrast to these previous findings, however, exposure to an acute stressor was necessary to uncover behavioral deficits in NodDKO mice, suggesting a different mechanism of action is involved in the response to immune stimulation. Increased serum corticosterone concentrations and decreased hippocampal GR expression observed in NodDKO(+WAS) suggests a unique role for NLR in maintaining HPA-axis signaling in response to acute stress.

Given the association between stress, hippocampal neuronal activation, and behavior (36, 48, 49), hippocampal neuronal activation was assessed by Arc expression and c-Fos activation. The increase in Arc expression that would otherwise be induced by stress was inhibited, and the numbers of c-Fos positive neurons were reduced in NodDKO(+WAS) mice. This indicates a lack of neuronal activation following exposure to WAS. Stress is also known to be detrimental for adult hippocampal neurogenesis (50), which has been shown to reduce memory recognition in rodents, as assessed by NOR tasks (30, 51). Similarly, both the number of immature neurons and hippocampal cell proliferation were impaired in NodDKO(+WAS) mice. Taken together, these findings demonstrate that behavioral impairments driven by NLR appear to involve alterations in hippocampal neurons, including activation and neurogenesis, which may lead to impaired consolidation of hippocampal-dependent memories and cognitive deficits.

Given the interaction between stress and hippocampal serotonergic signaling (52) we wanted to assess 5-HT biology in NodDKO mice. In the hippocampus, a lack of Nod1/Nod2 receptors resulted in significantly decreased 5-HT at both baseline and following exposure to acute stress compared to WT controls. This finding was associated with significantly reduced expression of both *Tph2* (the rate-limiting enzyme for neuronal 5-HT synthesis) and the heteroreceptor *5HTr1a* in NodDKO(+WAS) mice. While *Tph2* expression was not statistically reduced in NodDKO(-WAS) vs. WT(-WAS), significantly decreased hippocampal 5-HT concentrations in NodDKO(-WAS) mice compared to WT(-WAS) controls were detected. These data suggest that reduced *Tph2* expression may be sufficient to cause profound deficits in neurotransmitter production in the brain. Interestingly, both stress and elevated corticosterone levels have been shown to downregulate the expression and the functionality of *5HTr1a* through the alteration of *GR* expression in the hippocampus of both rats and subjects with a history of depression (53, 54). Highlighting a role for 5-HT in mediating cognitive function, administration of the selective post-synaptic *5HTr1a* agonist F15599 can ameliorate cognitive performance during the NOR task in a rat model of cognitive impairment (55). Moreover, pharmacological blockade of *5HTr1a* can reduce hippocampal neurogenesis in adult rats (56), and regulate memory (57) as well as emotional behavior (29). In addition to hippocampal changes, significantly lower 5-HT concentrations were observed in the brain stem of NodDKO(±WAS) mice, suggesting lower release of 5-HT to the hippocampus (35). Although reduced production of 5-HT was observed in both NodDKO(-WAS) and NodDKO(+WAS) mice, exposure to acute stress was necessary to uncover the behavioral deficits found in NodDKO(+WAS) mice. These data suggest a critical interaction between stress, NLR, and the 5-HT system in the regulation of memory and emotional behavior. Future studies characterizing the long-lasting effects of Nod1/Nod2 receptors on serotonergic signaling in response to chronic stress could support the concept that NLR play a crucial role in the pathophysiology of stress-related disorders, with implications for treatment.

To determine if the behavioral deficits observed in NodDKO(+WAS) mice were dependent on the impaired serotonergic system seen in the hippocampus, behavior and intestinal physiology were assessed in mice treated chronically with fluoxetine to increase 5-HT availability in the CNS. Restoration of cognitive impairment and reduced anxiety-and depressive-like behaviors were observed after chronic fluoxetine treatment, further supporting a role for 5-HT in mediating the behavioral deficits seen in NodDKO(+WAS) mice. In addition to increasing 5-HT concentration in the synaptic cleft by inhibiting SERT, the antidepressant effects of fluoxetine have been demonstrated to involve other mechanisms such as neurogenesis (32), increased BDNF levels (58), regulation of HPA-axis activity (59) and modulation of long-term potentiation (60). Therefore, while NLR appear to regulate serotonergic signaling in the brain, it remains to be determined at which point in the signaling pathway they are critical for maintaining 5-HT levels in the brain and periphery.

Since NLR are important in maintaining intestinal physiology (38) that can be detrimentally impacted by exposure to stress (61, 62), we assessed ileal and colonic mucosal barrier function in WT(±WAS) and NodDKO(±WAS) mice. While baseline ion transport (Isc) and permeability (G, FITC flux) were intact in NodDKO(-WAS), stress significantly increased Isc, G, and FITC flux in the ileum and/or the colon of NodDKO(+WAS) mice vs WT(+WAS) controls. These changes in GI physiology were associated with altered 5-HT signaling, with increases in *5HTr1a* and decreased *Tph1* expression seen in the ileum. Mucosal barrier deficits in NodDKO(+WAS) mice were not restored by fluoxetine administration, suggesting that fluoxetine ameliorates behavioral deficits via a mechanism independent of intestinal physiology. This may be due, in part, to the dual function of 5-HT in the GI tract, depending on whether it is released from enteric nerves or enteroendocrine cells (EC) in the mucosa (39), although future studies would be necessary to discern the specific contribution of each source in maintaining intestinal physiology when NLR are absent. Interestingly, activation of Nod1 has been shown to downregulate the activity and expression of SERT in epithelial Caco-2/T7 cells (63). Although these findings need to be confirmed *in vivo*, this suggests a potential interaction between NLR and the serotonergic system in the GI tract.

To assess if Nod1/2 deficient mice have reduced peripheral availability of Trp, the biochemical precursor for 5-HT, the concentration of this amino acid was quantified in the serum. The concentration of serum Trp was significantly reduced in NodDKO(+WAS) mice but could be restored by chronic fluoxetine treatment. Since CNS concentrations of Trp are predominantly determined by availability in the periphery (64), our data suggests a reduced supply of this amino acid precursor of 5-HT to the CNS as a potential mechanism by which peripheral Nod1/Nod2 receptors can influence the CNS.

Given our findings of altered physiology and 5-HT signaling in the GI tract in NodDKO mice, the role of IEC NLR expression in regulating gut-brain communication was assessed using cKO mice. Cognitive deficits and anxiety-like behavior were observed in VilCre^+^Nod1^f/f^(+WAS), but not VilCre^+^Nod2^f/f^(+WAS) mice, suggesting that Nod1 expression in IEC can regulate cognition and mood in response to stress. While the anxiety-like behavior parameters in our cKO mice did not precisely phenocopy the changes seen in NodDKO(+WAS) mice, the presence of anxiety-like behavior is consistent and supportive of a common behavioral feature between the two strains. While our findings highlight a crucial role for IEC Nod1 receptors in maintaining behavior, it also suggests a potential additional role of brain Nod1/2 receptors in the regulation of memory and emotional mood. Thus, the use of selective antagonists as well as cKO mice targeting Nod1/2 receptors in the brain would be valuable to further elucidate the role of NLR in regulating mood and cognition.

Although genetic knockout studies have revolutionized the understanding of how individual genes contribute to complex phenotypes such as behavior, environmental factors can also exert a strong influence. The influence of environmental vs. genetic factors can be assessed by co-housing two groups of animals of differing genotypes in a single cage. Using this approach, the transfer of gut-brain phenotypes has been observed ranging from WAS-induced enteritis in mice (65) to an anxiogenic phenotype transferred from a stressed strain of mice into a non-stressed strain (66). In our studies, co-housing WT and NodDKO mice failed to reverse the behavioral deficits found in NodDKO(+WAS). This suggests that the genetic deletion of NLR serves as the main determinant of the susceptibility to stress-induced alteration in mood and cognition seen in NodDKO(+WAS) mice. This finding is further supported by studies using our VilCre^+^Nod1^f/f^(+WAS) mice, which were littermates (and thus cage mates) of the VilCre^-^Nod1^f/f^(+WAS) mice that did not show behavioral alterations. Together, these findings highlight that the absence of Nod1 in IEC is responsible for maintaining behavioral deficits compared to WT controls.

In conclusion, Nod1/Nod2 receptors represent novel mediators of gut-brain axis signaling. Deficiencies in NLR signaling in mice causes increased susceptibility to stress-induced behavioral deficits, including cognitive function, anxiety-like behavior, and depressive-like behavior, as well as intestinal mucosal barrier defects. These effects were coupled with alterations in the serotonergic system that could be restored by chronic fluoxetine administration, suggesting a link between 5-HT signaling and NLR. Moreover, these findings also indicate that Nod1/Nod2 receptors may be novel targets for the treatment of stress-related gut-brain disorders.

## Acknowledgments

The authors would like to thank Mr Dirk Truman (Monash Gene Targeting Facility, Monash University) for designing the Nod1 targeting strategy and for generating the heterozygote animals. This research was supported by the NIH (1R01AT009365-01 and 5R21MH108154-01 to MGG) and by the NHMRC (Senior Research Fellowships APP1079904, 606476 and Project grants APP1011303, 1079930 to RF). Research at the Hudson Institute of Medical Research is supported by the Victorian Government’s Operational Infrastructure Support Program.

## Contributions

MMP, MGG, KEB and CR designed research; MMP performed animal behavior; MMP, PS, GR, KAW performed RNA extraction and qRT-PCRs; MMP, LRG, SEG, IBM, performed immunohistochemistry and confocal microscopy; MMP, GR and MB performed LC/MS analysis and data interpretation; CL provided the LC/MS equipment; MMP, CTF and MGG performed the Ussing chamber experiments; MXB and AJB provided the NodDKO breeders; RF, CM, and DJP designed the Nod1^f/f^ mice; MS performed pilot animal behavior experiments and pilot immunohistochemistry; MS, DJP, AJB, KEB and CR provided critical feedback on the manuscript; MMP and MGG analyzed and interpreted data; MMP and MGG wrote the paper.

## Competing interests

Authors declare no conflict of interest.

