## Supplementary figures and images for "Nod-like receptors are critical for gut-brain axis signaling"

### Supplemental Figure 1

#### Supplemental Figure 1 - Behavioral Deficits in NodDKO(+WAS) Mice are Independent from Sex

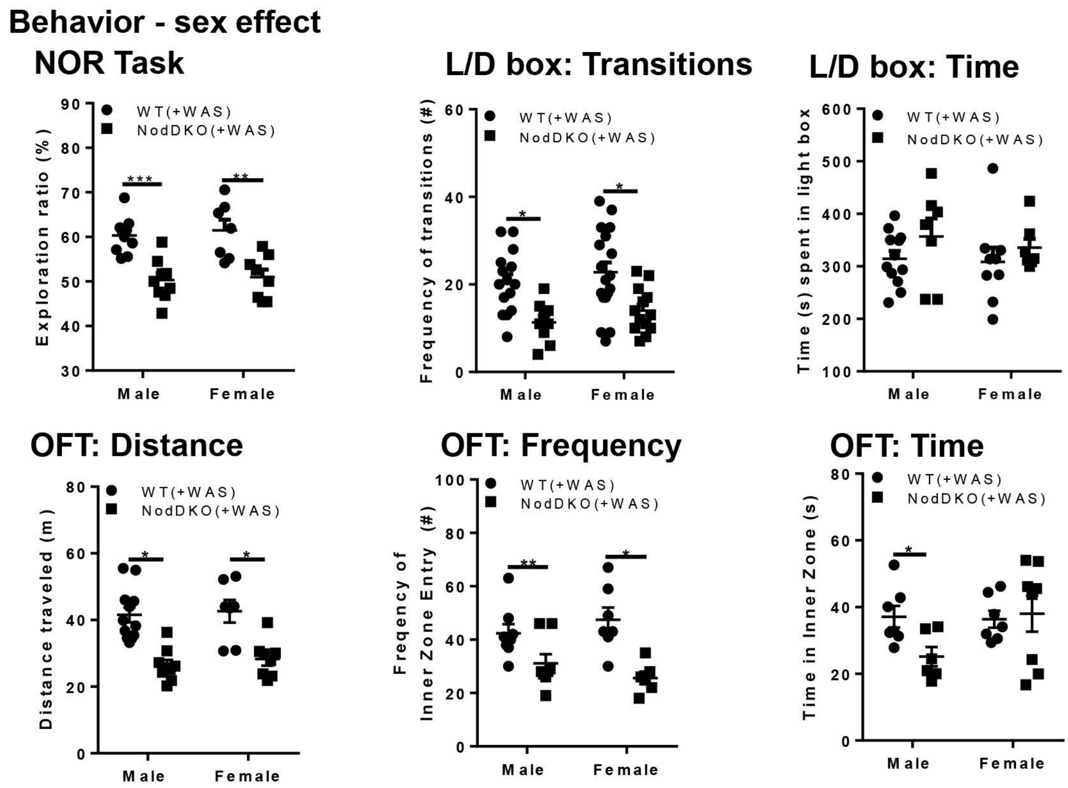

### Supplemental Figure 3

#### Supplemental Figure 3 - Basal Trp Serum levels are Lower in NodDKO(-WAS) Mice

##### Trp serum levels

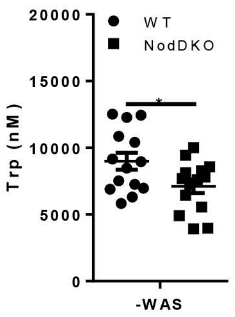
