## Supplemental Figure 2 for "Nod-like receptors are critical for gut-brain axis signaling"

### Supplemental Figure 2 - mRNA expression levels in the hippocampus and prefrontal cortex of NodDKO(+WAS) mice

#### A. Hippocampus

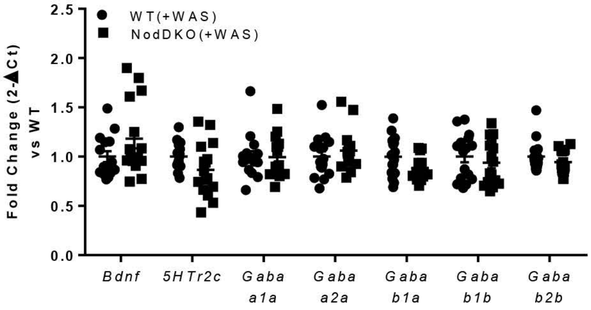


#### B. Prefrontal cortex

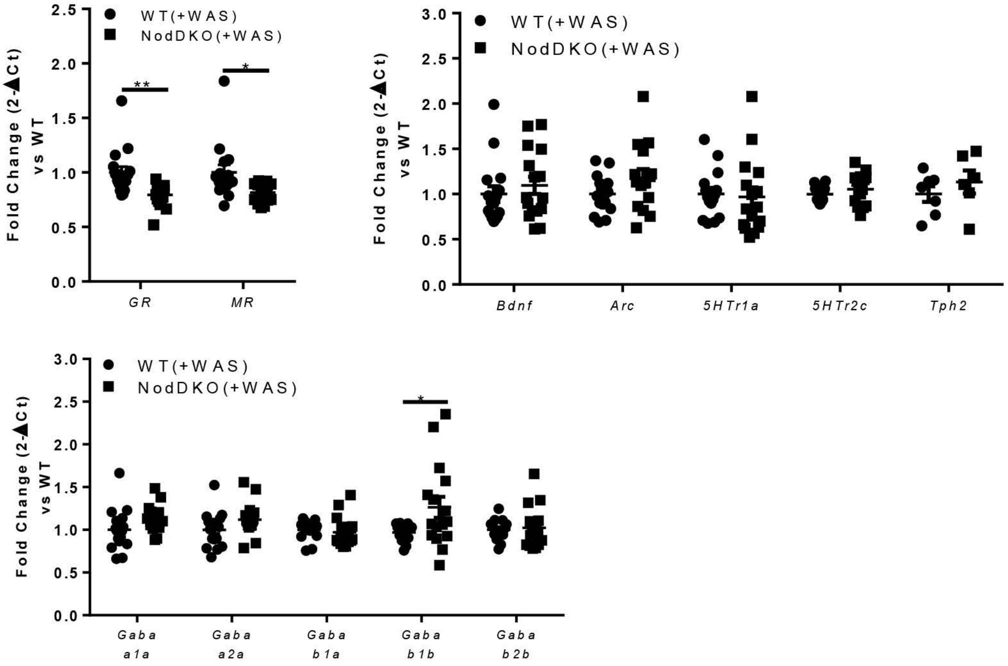
