## Supplemental Figure 4 for "Nod-like receptors are critical for gut-brain axis signaling"

**Supplemental Figure 4 - Chronic Fluoxetine Administration did not Restore Intestinal Physiology in NodDKO(+WAS) Mice**

### A. Ileum

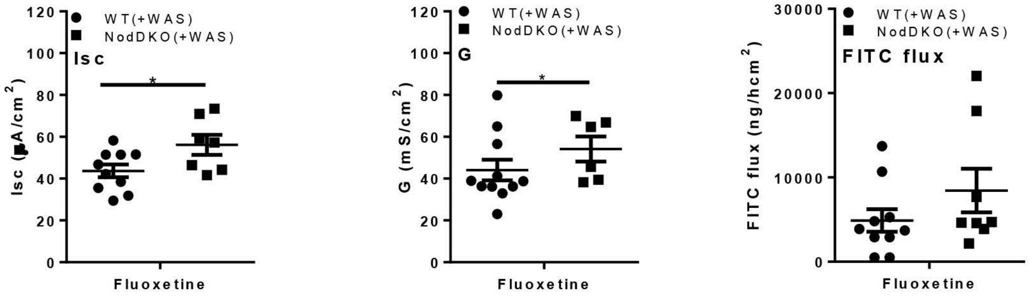


### B. Colon

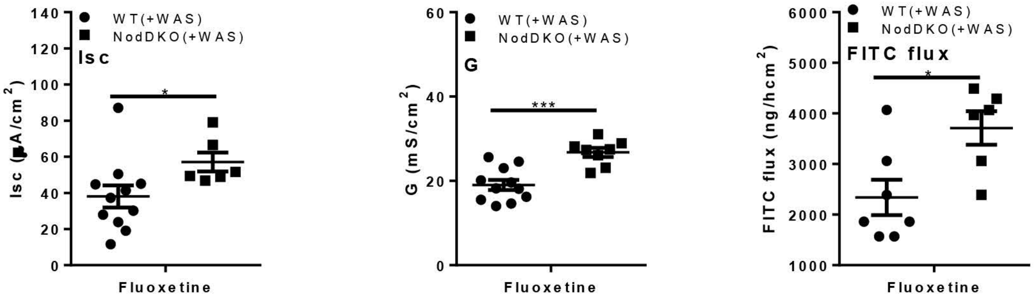
