## Supplemental Figure 5 for "Nod-like receptors are critical for gut-brain axis signaling"

**Supplemental Figure 5 - Behavioral Deficits are not Dependent on Intestinal Epithelial Nod2 Receptor**

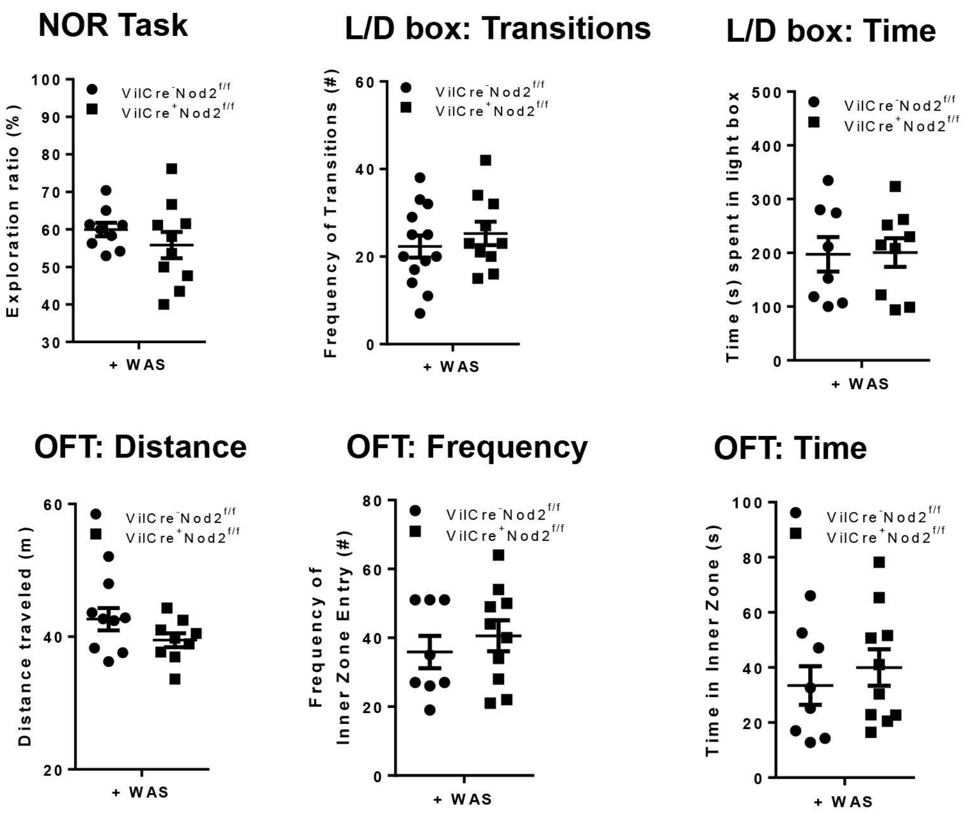
