## Supplemental Figure 6 for "Nod-like receptors are critical for gut-brain axis signaling"

**Supplemental Figure 6 - Schematic representing the floxed locus used for the generation of Nod1f/f mice**

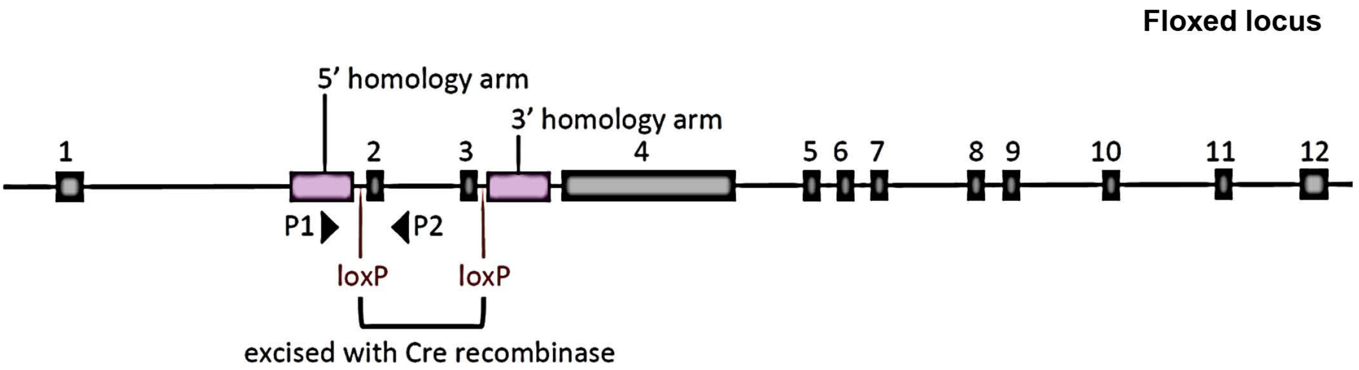
