## Supplemental Figure Legends for "Nod-like receptors are critical for gut-brain axis signaling"

**Supplemental Figure 1. Behavioral deficits in NodDKO(+WAS) mice are independent of sex.** Sex effect on NOR task, L/D box, and OFT (N = 7-18). Data are presented as mean ± SEM. (*P < 0.05; **P < 0.01, 2-way ANOVA).

**Supplemental Figure 2. mRNA expression levels in the hippocampus and prefrontal cortex of NodDKO(+WAS) mice.** (**A**) Hippocampus and (**B**) prefrontal cortex (PFC) qPCRs (N = 6-8). Data are presented as mean ± SEM. (*P < 0.05; **P < 0.01, unpaired Student’s t-test).

**Supplemental Figure 3. Basal Trp serum levels are lower in NodDKO(-WAS) mice.** Tryptophan (Trp) serum levels detected by LC/MS (N = 14). Data are presented as mean ± SEM. (*P < 0.05, unpaired Student’s t-test).

**Supplemental Figure 4**. **Chronic fluoxetine administration did not restore intestinal physiology in NodDKO(+WAS) mice.** (**A**) Ileum and (**B**) colon basal short circuit current [Isc], basal conductance [G] and FITC dextran flux assessment (N = 6-8). Data are presented as mean ± SEM. (*P < 0.05; **P < 0.01; ***P < 0.001, unpaired Student’s t-test).

**Supplemental Figure 5. Behavioral deficits are not dependent on intestinal epithelial Nod2 receptor.** Effect of intestinal Nod2 deletion on NOR task, L/D box, and OFT (N = 9-12). Data are presented as mean ± SEM (unpaired Student’s t-test).

**Supplemental Figure 6. Schematic representing the floxed locus for the generation of Nod1^f/f^ mice**
